# IndepthPathway: an integrated tool for in-depth pathway enrichment analysis based on bulk and single cell sequencing data

**DOI:** 10.1101/2022.08.28.505179

**Authors:** Sanghoon Lee, Letian Deng, Yue Wang, Kai Wang, Maureen A. Sartor, Xiaosong Wang

**Author notes:** **Corresponding Author**: Xiaosong Wang, M.D., Ph.D. Associate Professor of Pathology, UPMC Hillman Cancer Center, 5117 Centre Avenue, Pittsburgh, PA 15232.

## Abstract

Single-cell sequencing (SCS) enables exploring the pathways and processes of cells, and cell populations. However, there is a paucity of pathway enrichment methods designed to tolerate the high noise and low gene coverage of this technology. When gene expression data are noisy and signals are sparse, testing pathway enrichment based on the genes expression may not yield statistically significant results which is particularly problematic when detecting the pathways enriched in less abundant cells that are vulnerable to disturbances. In this project, we developed a Weighted Concept Signature Enrichment Analysis (WCSEA) algorithm specialized for pathway enrichment analysis from single cell transcriptomics (scRNA-seq). WCSEA took a broader approach for assessing the functional relations of pathway gene sets to differentially expressed genes, and leverage the cumulative signature of molecular concepts characteristic of the highly differentially expressed genes, which we termed as the universal concept signature, to tolerate the high noise and low coverage of this technology. We then incorporated WCSEA into a R package called “IndepthPathway” for biologists to broadly leverage this method for pathway analysis based on bulk and single cell sequencing data. Through simulating technical variability and dropouts in gene expression characteristic of scRNA-seq as well as benchmarking on a real dataset of matched single cell and bulk RNAseq data, we demonstrate that IndepthPathway presents outstanding stability and depth in pathway enrichment results under stochasticity of the data, thus will substantially improve the scientific rigor of the pathway analysis for single cell sequencing data. The IndepthPathway package is available through: https://github.com/wangxlab/IndepthPathway.

## BACKGROUND

The identification of causal pathways underlying physiological processes or disease initiation, progression, or therapeutic resistance based on proteomic, genomic, and transcriptomic datasets are major challenges of genomics research, particularly for single cell sequencing-based studies. Pathway Enrichment (PE) methods can be classified chronologically into three generations (Garcia-Campos, et al., 2015): 1) over-representation analysis (ORA) (Yu, et al., 2012) such as Fisher’s exact test, 2) functional class scoring such as Gene Set Enrichment Analysis (GSEA) (Subramanian, et al., 2005), and 3) network-topology based such as network enrichment analysis (NEA) (Jeggari, et al., 2018). Based on their dependency on gene weights (i.e., levels of differential expressions), PE methods can be classified into unweighted or weighted methods. The unweighted methods (i.e., ORA or NEA) are commonly used for analysis of genetic traits extracted from genomic analysis. For RNA-seq analysis, these methods can be used to identify the pathways within the user-defined DE genes based on thresholding. The weighted methods (i.e., GSEA) are commonly used for analysis of continuous expression data. Incorporating the degrees of differential gene expressions allows for detection of different magnitudes of pathway alterations.

It is notable that PE methods have distinct functionality from pathway activity inference, pathway and gene set over-dispersion analysis, regulon activity inference, or functional gene set inference. PE methods aim to test the enrichment of a wide array of functional pathways in differentially expressed genes thus are commonly used for pathway discovery. Whereas pathway activity inference tools assess the activity of handful pathways in each sample or single cell. Popular pathway activity inference tools such as PROGENy require consensus signatures obtained from publicly available perturbation experiments, which further limit the scope of the pathways to be assessed (Schubert, et al., 2018). Pathway and gene set over-dispersion analysis (i.e., PAGODA) is used to cluster cells into transcriptional subpopulations (Fan, et al., 2016). Regulon inference (i.e., AUCel) are designed for testing the activity of the large footprint regulons for gene regulatory network reconstruction (Aibar, et al., 2017), thus are not dedicated for pathways that have much smaller footprints. Functional gene set inference depends on PE methods for testing gene set enrichments based on matrix factorizations, which we will discuss further in the results section.

While there are a handful pathway activity inference methods for single cell sequencing (SCS) data, to date there is a paucity of PE methods tailored for SCS data despite its popularity for pathway discovery in biomedical research. Compared to bulk-sequencing, SCS enables exploring pathways and processes of individual cells. One of the most important analyses to achieve this is to identify the pathways that exhibit significant differences among different cell states or cell types. Due to low genome coverage and high amplification bias, bioinformatics tools developed for bulk sequencing may not work well with SCS data (Ning, et al., 2014; Poirion, et al., 2016). The conspicuous challenges of SCS data analysis include dissecting single cell expression variability, identifying rare cell populations, and understanding spatio-temporal transitions (Stegle, et al., 2015; Yuan, et al., 2017). In the case of PE analysis, the problem is less self-evident due to lack of golden standard and explicit modeling methods to assess the impact of SCS technical variability on the pathway interpretations. Further, it is even more challenging to detect the pathways enriched in less abundant cells which is more vulnerable to disturbances.

In our previous study, we sought to improve the reproducibility of PE analysis based on experimentally defined gene sets through leveraging the vast variety of knowledge-based gene sets including gene ontologies, pathways, interactions, and domains etc., to help inform complex functional relations, which we collectively termed as “molecular concepts”. Based on this idea, we developed a concept signature enrichment analysis (CSEA) for deep assessing the functional relations between pathway gene sets and a target gene list using a compendia of molecular concepts (Chi, et al., 2019). This method is grounded on the framework of shared concept signatures between gene sets, thus overcomes the limitations of the current algorithms relying on assessing gene set overlap. Here “a target gene list” is defined as a gene list of interest cataloged by comparative genomic analysis, and a “pathway gene set” is defined as a set of genes known to function in a certain pathway (**Fig. 1A**). The concept signatures of a target gene list are collectively termed as a Universal Concept Signature (uniConSig). CSEA deeply interprets the function of a target gene list via computing their overrepresentations in a wide array of molecular concepts, which were then used as weights to compute a cumulative genome-wide uniConSig score that represent the functional relevance of human genes underlying this target gene list (**Fig. 1D**). Then the UniConSig-sorted genome will be used for testing pathway enrichment through weighted Kolmogorov–Smirnov (WKS) tests (**Fig. 1E**). If a pathway gene set is functionally like the target gene list, the uniConSig scores will be high for that pathway genes, which will result in a high normalized enrichment score (NES) of that pathway. CSEA represents a new class of PE method that is distinct from all earlier generations with much improved depth in assessing gene set functional relations. A major advantage of CSEA over the approach based on interactome network topology is that CSEA is grounded on the framework of the vast knowledge databases, and thus can comprehensively assess most, if not all, functional aspects when computing the functional relations (Chi, et al., 2019). In this study, we further developed a weighted CSEA (WCSEA) method for pathway enrichment analysis based on a weighted gene list as in the function of GSEA. CSEA accept a list of target genes (i.e., up, or downregulated genes), whereas WCSEA accept a weighted target gene list (i.e., levels of differential expressions) as input data (**Supplementary Fig. 1**). Thus, CSEA interprets pathways in a gene set, whereas WCSEA interprets pathways in a weighted gene list.

**Figure 1.**
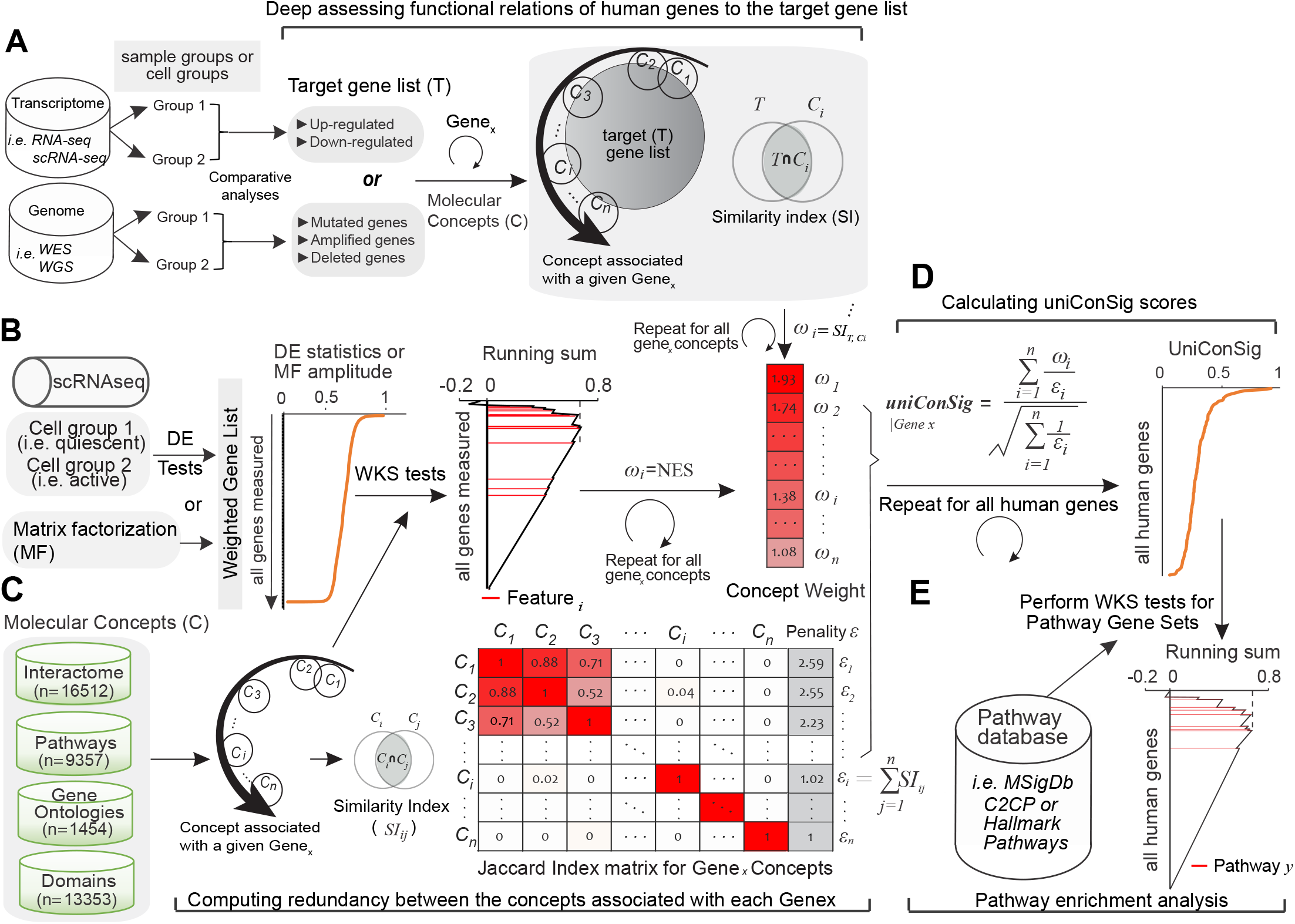
Schematic of the unweighted and weighted CSEA methods for PE analysis based on bulk or scRNA-seq data. (**A**) For CSEA analysis, a target gene list will be generated from genomic datasets (i.e., differentially expressed or mutated genes). CSEA achieves deep functional assessment of the target gene list by performing similarity tests to quantitate the enrichment of each molecular concept associated with a given Gene x in the target gene list. The resulting similarity index for each concept will be used for calculating uniConSig scores. (**B**) For WCSEA analysis, the weighted gene list can be created based on differential expression statistics comparing different cell groups, or from the amplitude matrix of a low dimensional component identified by matrix factorization methods. The uniConSig analysis achieves deep functional assessment of the weighted gene list (based on differential expression statistics) by performing Weighted KS tests to quantitate the enrichment of each molecular concept associated with a given Gene x in the highly weighted genes. The resulting NES score for each concept is used as weight for calculating the uniConSig score of Gene x, which will be normalized to the range of [0,1]. (**C**) These weights generated by similarity tests or WKS tests will be penalized by the degree of overlaps between the Gene x concepts, computed using their Ochiai’s similarity indexes based on molecular concept database. (**D**) The uniConSig scores will be computed using the indicated formula to reflect the functional relevance of human genes underlying the target gene list (for CSEA analysis) or highly weighted genes (for WCSEA analysis). (**E**) The enrichment of pathways in the pathway datasets (i.e., MSigDB C2CP canonical pathways, or hallmark pathways) in this uniConSig-sorted gene list will then be assessed using weighted K-S tests. The resulting normalized enrichment scores can be used as a quantitative measure of the functional associations between these pathways underlying the target gene list (CSEA) or weighted gene list (WCSEA).

We speculate that CSEA and WCSEA could be particularly suited for addressing the characteristics of SCS data. The rationale is that CSEA and WCSEA may tolerate the high noise and stochasticity of SCS data via leveraging the stable concept signatures of the target gene list to improve the reproducibility and scientific rigor of PE results. While the expressions of individual genes may be noisy, coordinated upregulation of genes within a gene set could provide a more stable signature. A similar principle has been leveraged by PAGODA to cluster cells into transcriptional subpopulations (Fan, et al., 2016). We then further develop a purpose-built PE package called “IndepthPathway” for deep pathway enrichment analysis from bulk and single cell sequencing data that took a broader approach for assessing gene set relations and leverage the universal concept signature of the target gene list to tolerate the high noise and low gene coverage of this technology. Our results show that IndepthPathway achieved better reproducibility under technical noise simulations of single cell sequencing data as well as in real matched bulk and single cell RNAseq data compared to other popular PE methods. Empowered by its strength in deep functional interpretation, IndepthPathway will have broad applications on inferring the pathway alterations from bulk and SCS data.

## METHODS

### Compiling molecular concept and pathway datasets

The uniConSig score calculation and deep functional interpretation for a target gene list rely on the precompiled human molecular concept databases. To generate a comprehensive knowledge base, we compiled 45,522 molecular concepts from the Molecular Signatures Database (MSigDB) C2CP pathways, or hallmark (v.7.0, https://www.gsea-msigdb.org/gsea/msigdb) (Zhao, et al., 2017), NCBI EntrezGene interactome (Brown, et al., 2015) database, the Pathway commons database (Rodchenkov, et al., 2020) (http://www.pathwaycommons.org), and conserved domain database (Brown, et al., 2015).

### Weighted concept signature enrichment analysis for pathway interpretation

To generate target gene lists, users can perform differential expression analysis of single-cell transcriptomic data based on their preferred methods such as Limma or Single Cell Differential Expression (SCDE) (Kharchenko, et al., 2014). The identified up- or downregulated gene lists were used as the target gene list for CSEA analysis. For WCSEA analysis, we used the signed statistical significance of the expression difference for the target genes as weights for uniConSig score calculation for genes and for pathway enrichment analysis.

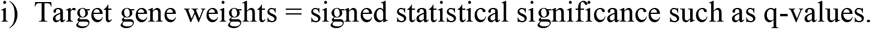

The basic algorithm to calculate uniConSig score for an experimental gene list in CSEA analysis is explained in our previous study (Chi, et al., 2020). The uniConSig scores assess the functional associations of human genes underlying the experimental gene list based on the similarities between the experimental gene list and the molecular concepts associated with each Gene_*x*_ in the genome. In WCSEA analysis, the calculation of uniConSig scores from a weighted gene list is based on the enrichment of each molecular concepts associated with Gene *x* in the weighted gene list calculated using weighted K-S test. The resulting normalized enrichment score (NES) for each concept is used as a weight *ω_i_* for calculating the genome-wide uniConSig score.

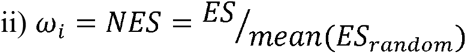

To reduce the inflation effects of redundant concepts on the uniConSig scores, we penalized the scores based on the similarity matrix of molecular concepts associated with each Gene_*x*_ in the genome. We generated the similarity matrix by computing the Otsuka-Ochiai coefficient between molecular concepts.

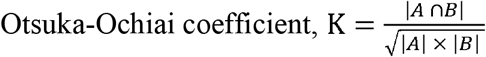

Compared to the Jaccard Index used in our previous study (Chi, et al., 2019), the Otsuka-Ochiai coefficient can better assess the similarity between a small molecular concept and a large molecular concept. To demonstrate this, we identified 300 large and small concept pairs in which each large concept contains all genes in the paired small concept, and calculated similarity indexes by both methods (**Supplementary Fig. 2**). Otsuka-Ochiai coefficient produced much higher similarity scores for these 100% overlapping concepting pairs than Jaccard Index. We then introduced a penalization factor (*ε*) calculated for each molecular concept *i* based on the cumulated similarities of concept *i* with other molecular concepts associated with Gene_*x*_. The Effective Weight (EW) is then calculated as:

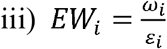

Finally, the uniConSig score of the given Gene_x_ is calculated as following:

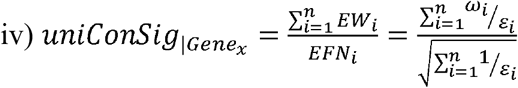

The cumulative genome-wide uniConSig scores reflect the functional relevance of the human genes underlying the highly weighted target genes, which can be interpreted as the adjusted average of normalized *ω_i_*. The normalized enrichment scores of pathway gene sets were then calculated by the weighted step-up of the random walk K-S test using uniConSig sorted human gene list to assess the functional associations of these pathways with the original weighted gene list. As in GSEA, users can interpret the most important pathways based on their ranks.

### Disambiguation of the crosstalk effects between similar pathways

Many pathways have varying degrees of overlap each other due to their shared genes. To reveal independent pathway modules, here we implemented disambiguation steps to correct the crosstalk effects between the enriched pathways, as previously described (Donato, et al., 2013). The enriched pathways should be independently enriched even after deleting the genes overlapped with the other similar pathway. Users can choose Bonferroni or Benjamini-Hochberg (BH) method for multiple testing correction of the p values. In our study the significance of the disambiguation process was determined by adjusted p-value < 0.01 based on “Bonferroni” method. For visualization of pathway association network, we removed the overlapping pathways in the top 30 pathways of WCSEA or GSEA output. We then assessed the functional association between the disambiguated pathways using our CSEA method as we described in our previous study(Chi, et al., 2020). The associations of the top pathways were visualized using either association heatmap or the pathway network. We used the corrplot R package (ver. 0.92) for the association heatmap and the igraph R package (ver. 1.2.4.2) for pathway network.

### Pathway enrichment analysis based on three scRNAseq datasets

We applied CSEA, WCSEA, GSEA, and other PE tools to analyze three public scRNA-seq datasets. The first dataset is GSE68981 that compared the active and quiescent populations of hematopoietic stem cells (HSCs, n=112) (Yang, et al., 2017). The second scRNAseq dataset is GSE75748 that profiled four lineage-specific progenitor cells derived from human embryonic stem cells (hESCs) (Chu, et al., 2016). More important, this dataset provide the matched bulk RNA-seq for hESC cells (n=4) and derived endothelial cells (EC, n=3) cells, which can be used for directly comparing the PE results based on real matched bulk and single cell RNAseq data. We thus performed PE analysis comparing hESC cells with EC cells using both bulk and scRNAseq data (212 hESC cells vs 105 ES cells). The third scRNA-seq dataset is GSE132172 that contains scRNA-seq data for human neural stem cells (n=59) and glioblastoma (GBM) cancer stem cells (n=75) (Zhao, et al., 2019). PE analyses were performed to compare the aforementioned cell types using multiple PE tools.

### Simulations of multiple faceted variability of scRNA-seq and dropouts in gene expression

To benchmark the performance of PE tools, we simulated the multi-faceted variability of scRNA-seq data using Synthetic model of multiple variability factors for Simulation (SymSim) (Zhang, et al., 2019). We simulated technical variations by noise intrinsic to the process of transcription, extrinsic indication of different cell states, and low sensitivity in measurement. The example code to simulate scRNA-seq data is included in our IndepthPathway R package. In addition, we simulated dropouts in gene expressions by randomly deleting 20% or 50% of the expressed genes. We then compared the performance of CSEA, WCSEA and other PE discovery tools, such as Over-Representation Analysis (ORA) (Yu, et al., 2012), Network Enrichment Assessment (NEA) (Jeggari, et al., 2018), Gene Set Enrichment Analysis (GSEA) (Subramanian, et al., 2005), UniPath (Chawla, et al., 2021), and iDEA (Ma, et al., 2020) based on the variations in pathway ranks under simulated technical variations. ORA is implemented based on R package clusterProfiler (ver. 4.6.0), and NEA is implemented based on NEArender R package (ver. 1.5). UniPath was implemented using GitHub R package (ver, 0.1.0, https://reggenlab.github.io/UniPathWeb/). iDEA was implemented using GitHub R package (https://xzhoulab.github.io/iDEA/installation/).

### Visualization of enriched pathways from IndepthPathway analysis

Here we provide five modules for visualization of enriched pathways. The first option generates a heatmap revealing the functional association between the top up- or down-regulated pathways (**Supplementary Fig. 3**). The second option generate a network revealing the functional associations of the top up or downregulated pathways. The third option generates an enrichment plot for one selected pathway as in GSEA (**Supplementary Fig. 4A**). The fourth option generates a heatmap to represent differentially expressed genes from a specific pathway gene set as in GSEA (**Supplementary Fig. 4B**). The fifth option generates an interactome network that visualize the differential expression levels of genes in a selected pathway and their interactions with a target gene of interest based on human reference binary protein interactome (Luck, et al., 2020) and Entrez gene reference interactome database (Maglott, et al., 2011) (**Supplementary Fig. 5**).

## RESULTS

### Development of a weighted CSEA method to enhance its utility and reproducibility in detecting pathway alterations from single cell transcriptomics data

Next, we developed a Weighted CSEA (WCSEA) method to enhance its functionality and reproducibility in analysis of single cell transcriptomics. We propose WCSEA to achieve deep functional assessment of the weighted gene list (based on differential expression statistics) by performing Weighted KS tests to quantitate the enrichment of each molecular concept associated with a given Gene_*x*_ in the highly weighted genes. The resulting NES score for each concept is used as weight for calculating the uniConSig score of Gene_*x*_ (**Fig. 1B**). These weights are then penalized by the degree of overlaps between the Gene_*x*_ concepts, computed based on their Ochiai’s similarity indexes (**Fig. 1C**). The resulting uniConSig scores reflect the functional relevance of human genes underlying the highly weighted genes in the list (i.e., highly overexpressed genes) (**Fig. 1D**). Then pathway enrichment analysis will be performed on this genome-wide weighted gene list based on uniConSig scores (**Fig. 1E**). WCSEA leverages the stable concept signatures of highly weighted genes to create a buffer zone for tolerating the high noise and stochasticity of scRNA-seq data.

### Development of IndepthPathway package for biologists to broadly leverage this technology to enhance pathway discovery from experimentally defined gene sets and SCS data

Next, we implemented CSEA and WCSEA in a R package called “IndepthPathway” to widely disseminate this technology. This purpose-built tool allows users to provide a weighted or unweighted target gene list defined by comparative analysis of bulk or single cell sequencing data for WCSEA or CSEA analysis respectively (Supplementary Fig. 1). Users can perform differential expression (DE) analysis between the comparing sample groups or cell groups using the bulk or single cell sequencing DE methods, such as DESeq2 (Love et al. 2014 Genome Biology), Limma (Ritchie et al. 2015 NAR), and SCDE (Kharchenko, et al., 2014) and the resulting statistical values of user’s choice can be used to generate a weighted gene list for WCSEA analysis. This will allow user to implement their preferred DE methods to identify differentially expressed genes, and we have implemented the Limma and SCDE methods in our package. Over- or under-expressed pathways should be analyzed separately. Users can also implement unsupervised Matrix factorizations (MF) techniques to reveal the low-dimensional components (LDC) for functional pathway inference (Stein-O’Brien, et al., 2018). MF is not a PE method, but instead, it relies on PE analysis to explore the pathways enriched in an LDC using its amplitude matrix which is known as functional pathway inference. Users can select popular MF methods such as UMAP (Becht, et al., 2018) or t-SNE (Linderman, et al., 2019), and perform WCSEA analysis for selected LDC to identify the key pathways underlying that component.

For deep functional interpretations, we have precompiled the human concept datasets from the molecular signatures database (MSigDB) (Subramanian, et al., 2005), NCBI EntrezGene interactome database (Brown, et al., 2015), and conserved domain database (Brown, et al., 2015). Users can use our pre-compiled human concept datasets or provide their own concept dataset. We expect inclusion of the more available concepts will substantially “thicken” the “buffering zone” via increasing the number of signature concepts, thus will further improve the stability of WCSEA under technical noise. For pathway enrichment analysis, the pathway gene sets for curated canonical pathways and hallmark pathways have been precompiled from MSigDB (Subramanian, et al., 2005) and provided to users. Users may select the pre-compiled pathway gene sets or provide their own pathway gene sets in the Gene Matrix Transposed file format (*.gmt). In addition, we have integrated downstream analysis modules to improve its usefulness. Following pathway enrichment analysis, we have implemented a module for users to disambiguate and identify independent pathway modules via correcting the crosstalk effects between pathways, as previously reported (Donato, et al., 2013). The pathway enrichment results include the pathways ranked by NES scores, together with the p-values and FDR q-values. We also implemented a module to allow users to generate a pathway network showing the functional relations between significant pathways which are computed based on the CSEA method, and a module to visualize the differential expression and interactome of core enriched genes in a selected pathway related to a select target gene of interest, which is a powerful tool to explore the mechanisms.

### IndepthPathway provided more in-depth view of complex pathway alternations from scRNAseq data

Next, we applied the IndepthPathway tool to examine the pathways enriched in active vs quiescent populations of hematopoietic stem cells (HSC) based on a publishes single cell sequencing dataset (Yang, et al., 2017). Differential gene expressions between active and quiescent HSC are compared and the resulting statistical values are used as weights for WCSEA analysis. Following WCSEA analysis, the top 30 upregulated and top 30 downregulated pathways were disambiguated as explained in the Methods section and plotted into a network space based on the functional associations of these pathways computed through WCSEA analysis (**Fig. 2A**). Our results show that the WCSEA methods can comprehensively detect the pathways known to characterize active HSC: 1) Upregulation of E2F targets, DNA replication, and G2M checkpoint pathways, indicating increased cell cycle activities (Kwon, et al., 2017; Yang, et al., 2017); 2) Upregulation of MYC targets known to control HSC self-renewal and differentiation (Wilson, et al., 2004); 3) Upregulation of spliceosome, which is consistent with the increased total RNA amount in active HSCs (Huttmann, et al., 2001); 4) Downregulation of pathway required for the quiescent HSCs to respond to interferon α/β stimulation (Essers, et al., 2009; Robb, 2007); 5) upregulation of various DNA repair pathways. It is reported that DNA damages accumulated during quiescent state will be repaired upon entry into active state (Beerman, et al., 2014). In contrast, the commonly used Gene Set Enrichment Analysis (GSEA) (Subramanian, et al., 2005) appears to enrich the most significant pathways but lack the depth to interpret complex pathway alternations (**Fig. 2B**). Furthermore, interrogation of interactome network revealed the direct role of MYC in regulating the replicative complex in addition to its function in transcriptional regulation of cell cycle (**Supplementary Fig. 5**), consistent with previous report (Dominguez-Sola, et al., 2007).

**Figure 2.**
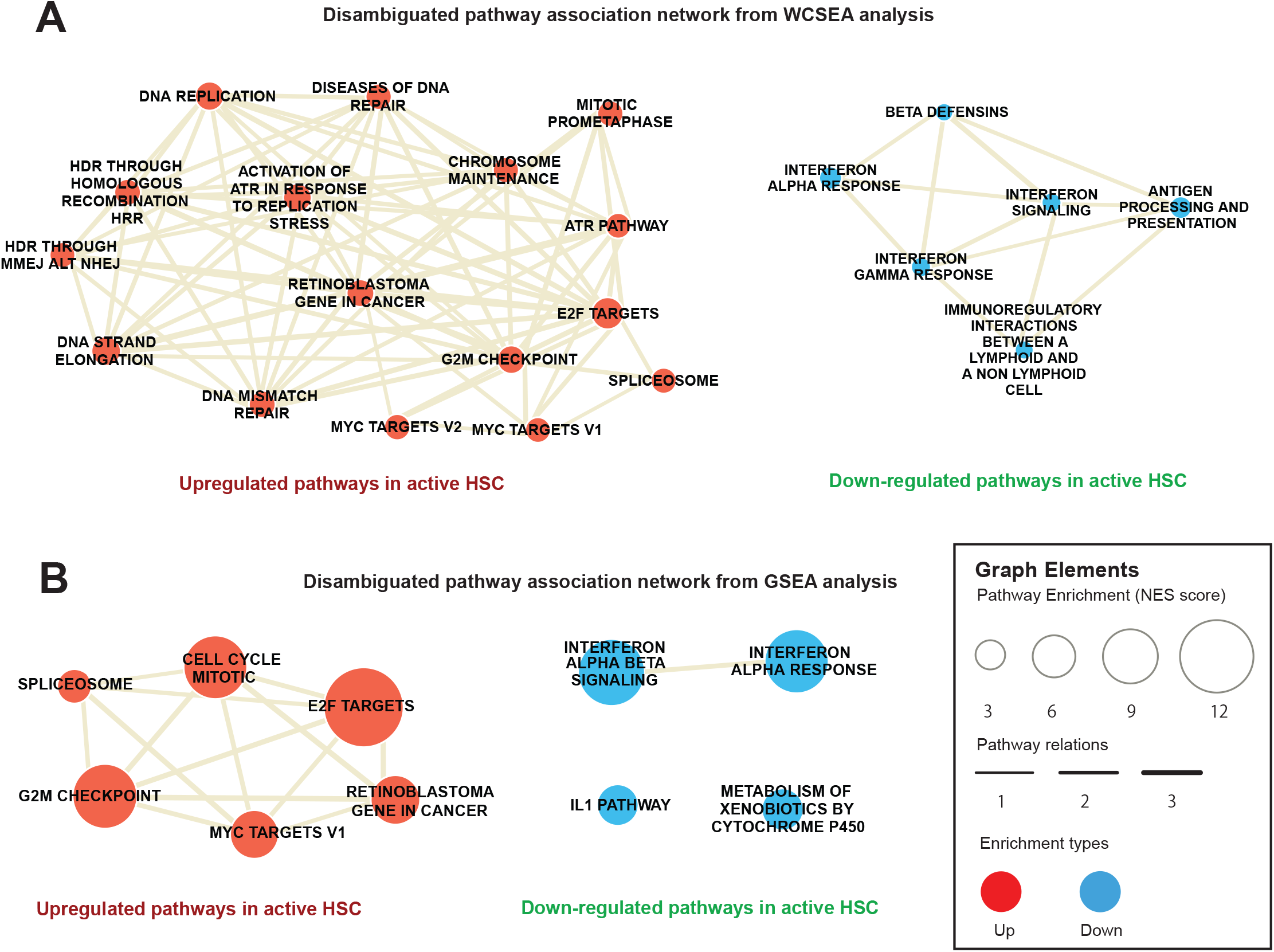
Pathway characteristics of active HSCs revealed by WCSEA or GSEA based on a single cell sequencing dataset. Pathway association networks showing the top upregulated and downregulated pathways in active HSCs compared to quiescent HSCs detected by WCSEA (**A**) or GSEA (**B**). Top 30 upregulated and top 30 downregulated pathways based on WCSEA or GSEA analysis are disambiguated, and the resulting non-ambiguous pathways are plotted in the network. The node size correlates with the levels of enrichments of the respective pathways, and the edge thickness correlates with the levels of functional associations between the pathways.

We then compared IndepthPathway with other PE methods in detecting significantly enriched pathways in active vs quiescent populations of HSC. We included representative PE tools of each generations including the GSEA, NEA, ORA, among which GSEA accepts weighted gene list based on DGE as in WCSEA, whereas NEA and ORA accept up or downregulated gene sets as in CSEA. In addition, we also included two pathway tools specialized for scRNAseq data, including a new PE tool called iDEA that carries out integrative differential expression and gene set enrichment analysis (Ma, et al., 2020), and a popular pathway activity inference tool called uniPath (Chawla, et al., 2021). As discussed above, pathway activity inference tools produce activity scores for each pathway in each single cell and additional statistical analysis will be required for PE analysis to compare the pathway activities between the different cell groups, and we have implemented in our analysis. We then constructed kernel density of p-values calculated for enriched pathways detected by different PE tools to view the distribution of p-values within 0.05 significance range (**Fig. 3A**). The left skewed density of the p-values for the detected pathways indicate that WCSEA identified more significantly enriched pathways than other PE methods. We also counted the number of significant pathways with increasing FDR q-value cutoffs for different PE methods (**Fig. 3B**). CSEA and WCSEA identified more enriched pathways with low false positive rates compared to other PE methods.

**Figure 3.**
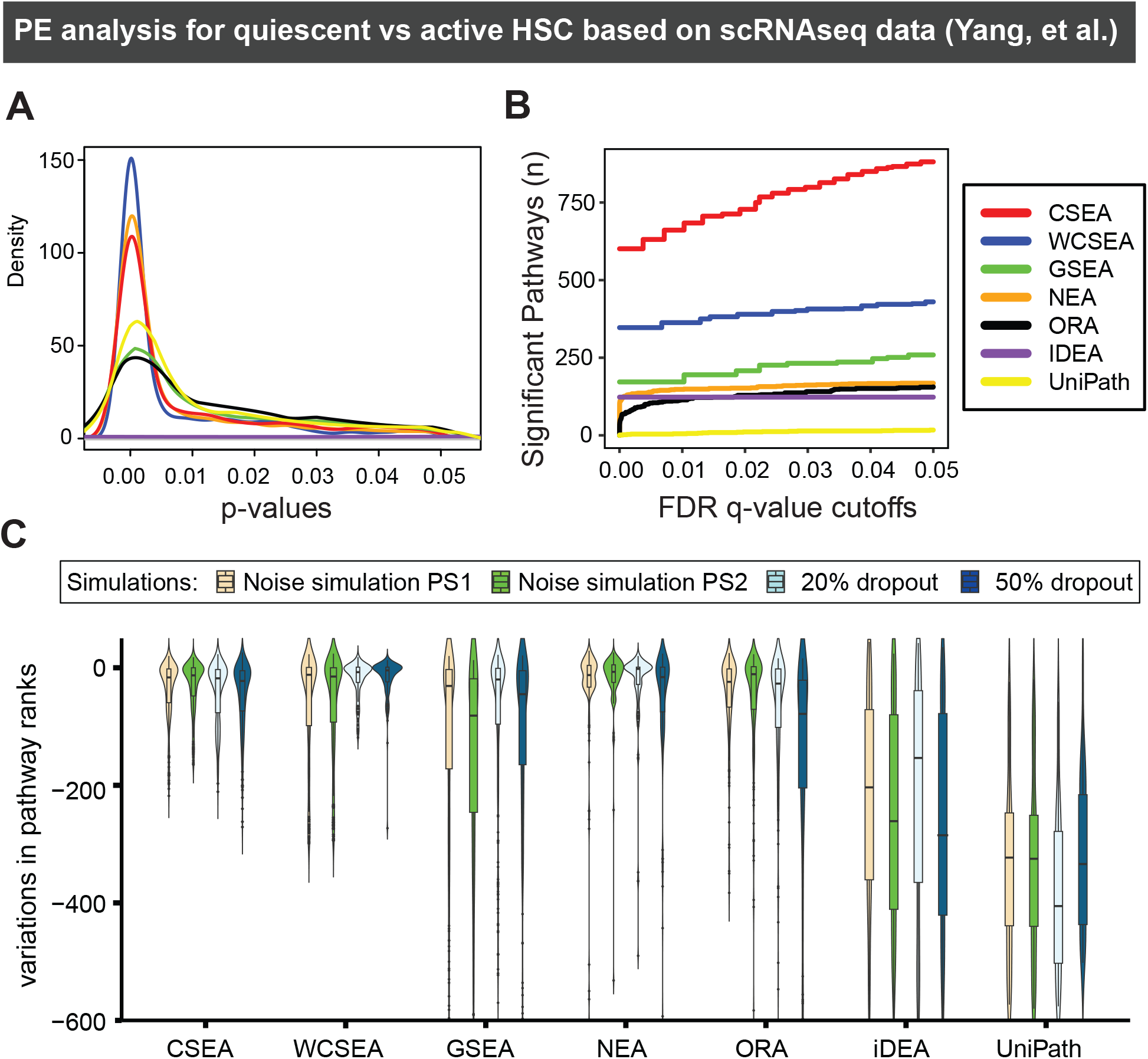
The performance of CSEA, WCSEA, and other PE tools on detecting pathways characteristic of active HSC based on scRNA-seq dataset and the reproducibility under simulated technical variability and dropouts in gene expression. (**A**) Kernal density of p-values from CSEA, WCSEA, and other PE tools. The density plot display the significantly enriched pathways detected by each PE tool within p-value 0.05. (**B**) The number of identified enriched pathways by each PE tools based on increasing FDR q-value cutoffs from 1×10^-05^ up to 0.05. **(C)** Noise simulations are performed using SymSim with two sets of parameters representing different noise levels. The dropouts are simulated via randomly deleting 20% or 50% expressed genes. Five permutations are done for each simulation. Read counts for 177 HSC cells (77 quiescent HSCs and 100 active HSCs) are used as true counts before simulations. SCDE is used for DE analysis, and the results are used for PE analysis. Variations in pathway ranks for top 30 up- or down-regulated pathways in quiescent HSCs are used for benchmarking with negative variations indicating reduced significance of the pathways.

### IndepthPathway yielded reproducible pathway results under simulations of multiple faceted variability of scRNA-seq and dropouts in gene expression

Next, we sought to examine the performance of IndepthPathway under the simulated technical variability of scRNA-seq data. This can be achieved through “SymSim” that explicitly models the multiple faceted variability of single cell RNA-seq (scRNA-seq) experiments (Zhang, et al., 2019). In addition, we simulated dropouts in gene expressions by randomly deleting 20% or 50% of expressed genes and assessed the changes in pathway ranks from the original results. We then tested the performance of IndepthPathway on pathway discovery from scRNA-seq with or without simulated technical variabilities (**Fig. 3C**). Pathways are usually interpreted based on their enrichment ranks, thus the rank variations for the top-30 up- and down-regulated pathways identified by these methods compared to the original PE results are used for benchmarking.

As expected, the simulations resulted in high variations in the pathway ranks from GSEA, NEA, ORA, iDEA and UniPath (**Fig. 3C**). This suggests that PE tools developed for bulk RNA-seq or even for scRNA-seq may yield inconsistent results under technical variability of scRNA-seq. When gene expression data are noisy and signals are sparse, testing pathway enrichment based on the genes measured may not yield statistically significant results. Tolerating dropouts in gene expression will require deep interpreting the functions of the differently expressed genes in the comparing pathways, instead of simply mapping the gene names in the pathways. Indeed, our result showed that CSEA and WCSEA produced overall lowest variations in pathway ranks with highest density around zero compared to other PE methods under simulations of both technical noises and gene expression dropouts. Together, this result supports the advantage of IndepthPathway on reproducibility under high technical noise and dropouts in gene expression.

### IndepthPathway results show outstanding reproducibility in matched bulk and single cell RNAseq data, offering in-depth view of pathway alterations in two additional scRNAseq datasets

Next, we sought to show the variability of PE results using the real dataset of matched bulk RNAseq and scRNAseq on the same cell line models. Through extensive literature investigation, we identified a dataset, GSE75748, with matched bulk RNAseq and scRNA-seq of Human Embryonic Stem Cells (hESCs) and derived endothelial cells (EC). We thus performed pathway enrichment analysis for EC vs hESC using different PE tools. In terms of kernel density of p-values for enriched pathways and significant pathways under different FDR q-value cutoffs, CSEA and WCSEA showed outstanding performance compared to other PE methods (**Fig. 4A-B**). We then examined variations of pathway ranks between PE results based on the bulk RNA-seq and scRNA-seq datasets which is a most important test of variability of PE tools (**Fig. 4C**). Our results show that CSEA and WCSA outperformed all other PE tools we tested, including GSEA, NEA, ORA, iDEA, and uniPath, showing much less variations in PE results. Likewise, in the simulations of noise or dropouts, CSEA and WCSEA showed much lower variations in pathway ranks than other PE tools (**Supplementary Fig. 6**).

**Figure 4.**
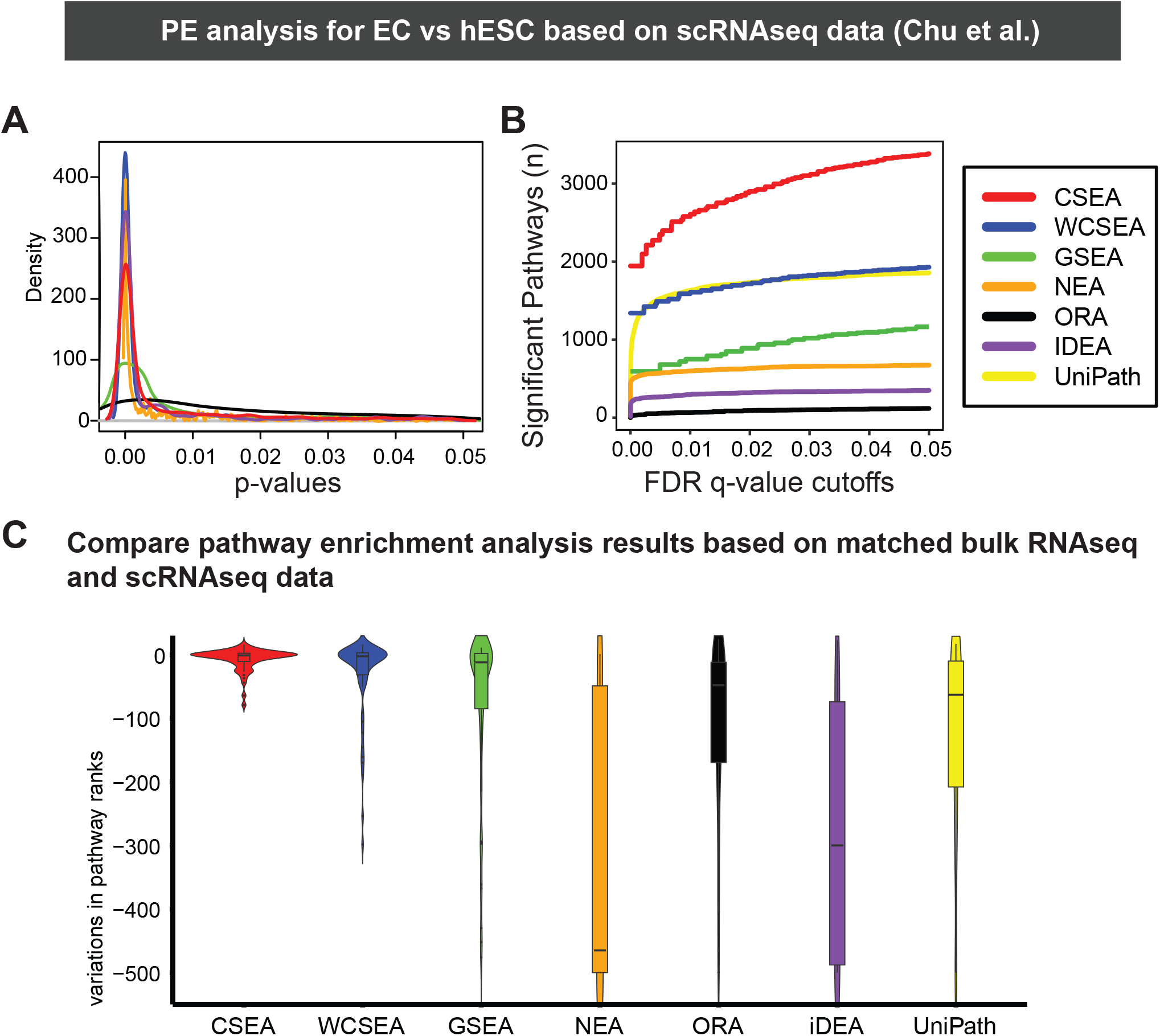
The performance and reproducibility of CSEA, WCSEA, and other PE tools on detecting pathways characteristic of endothelial cells compared to embryonic stem cells based on matched bulk and scRNA-seq dataset. (**A**) Kernal density plots of p-values from CSEA, WCSEA, and other PE tools. (**B**) The number of identified enriched pathways by each PE tools based on increasing FDR q-value from 1×10^-5^ up to 0.05. (**C**) The reproducibility of CSEA, WCSEA, and other PE tools in real matched bulk and single cell sequencing datasets comparing embryonic stem cells and derived endothelial cells. Variations of the pathway ranks between bulk RNA-seq and scRNA-seq datasets are shown in the figure with negative variations indicating reduced significance of the pathways. The matched bulk and single cell RNAseq data are from GSE75748.

Furthermore, WCSEA offered more in-depth and meaningful view of the pathway alterations during endothelial differentiation than GSEA, particularly for upregulated pathways (**Fig. 5**). For example, WCSEA was able to comprehensively detect key pathways in endothelial differentiation and function, such as angiogenesis, PDGFRB (Rolny, et al., 2006), and VEGF pathways (Li, et al., 2017), integrin (Toya, et al., 2015), FAK, and RhoGTPases pathways (Shen, et al., 2013), IL6 (Fan, et al., 2008), EPHA2, and inflammatory pathways (Funk, et al., 2012). These results suggest that the high noise and low gene coverage of scRNAseq data indeed can result in high variations in PE analysis and support the utility of IndepthPathway to overcome this issue.

**Figure 5.**
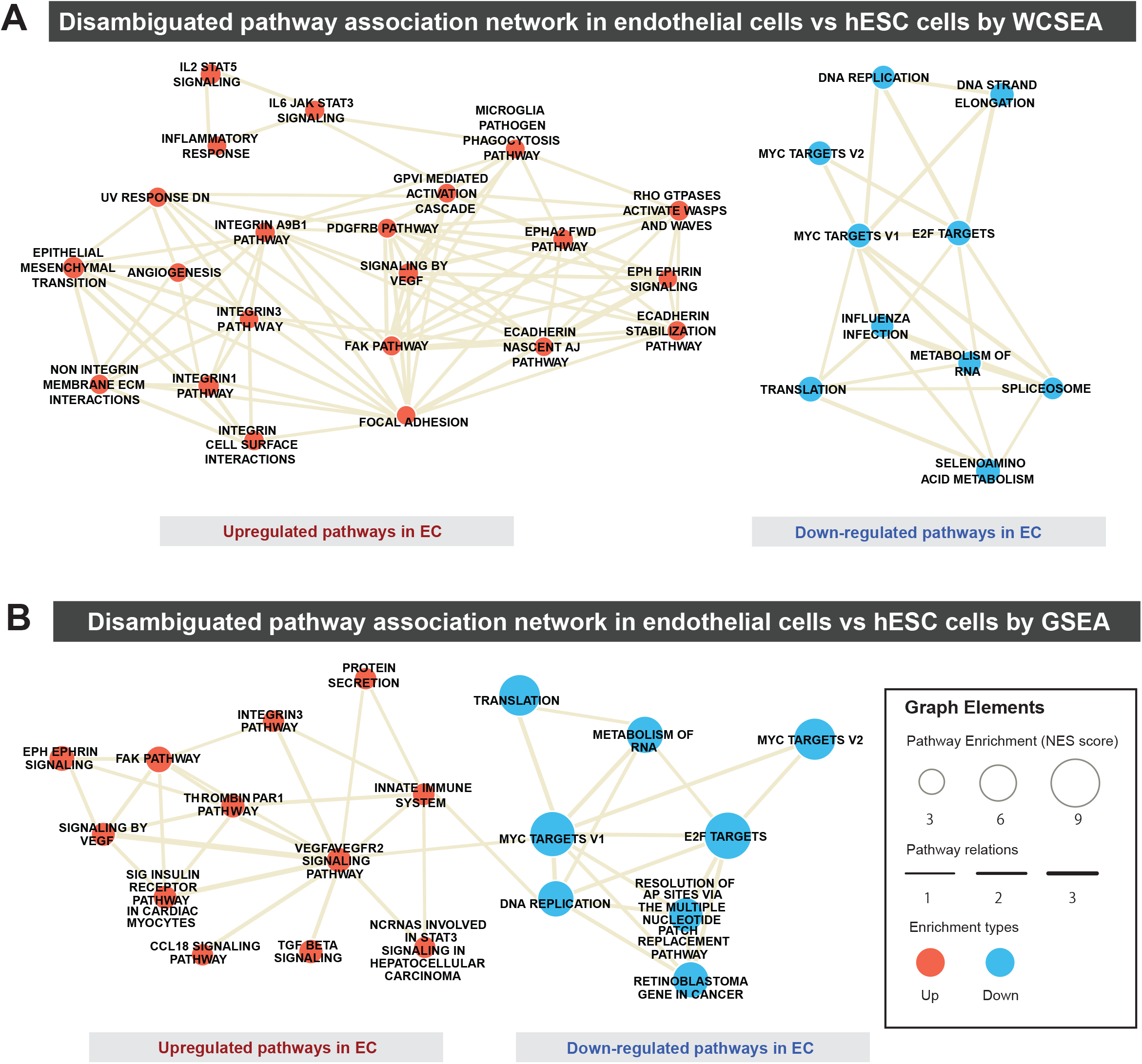
Pathways characteristics of endothelial cells differentiated from embryonic stem cells revealed by WCSEA or GSEA. Pathway association networks showing the top upregulated and downregulated pathways in endothelial cells compared to H1 hESC cells detected by WCSEA (**A**) or GSEA (**B**). Top 30 upregulated and top 30 downregulated pathways based on WCSEA or GSEA analysis are disambiguated, and the resulting non-ambiguous pathways are plotted in the network. The node size correlates with the levels of enrichments of the respective pathways, and the edge thickness correlates with the levels of functional associations between the pathways.

Moreover, we also analyzed a third dataset, GSE132172, that compared neural stem cell with GBM stem cells. To test if CSEA/WCSEA can reveal more significant pathways from the scRNA-seq data, we also compared the density of p-values and the number of significant pathways at different FDR cutoffs. We show CSEA and WCSEA revealed more significant pathways than all other tools (**Supplementary Fig. 7A-B**), providing in-depth view of the enriched pathways (**Supplementary Fig. 7C-D**).

## DISCUSSION

Bioinformatics tools for bulk-sequencing may not work well with SCS data due to the low genome coverage and high amplification bias. In case of pathway enrichment (PE) analysis, the problem is less self-evident and suitability of existing PE methods on SCS data are hard to assess due to the lack of golden standard. When gene expression data are noisy and signals are sparse, testing pathway enrichment (PE) based on the genes measured may not yield statistically significant results. Indeed, our analyses revealed that popular PE methods yielded highly inconsistent results under the simulated multiple faceted variability of single cell RNA-seq, and when directly comparing real matched scRNAseq and bulk RNAseq data. Here we introduce a new weighted CSEA analysis for deep assessing the pathways enriched in a weighted gene list such as differentially expressions through exploiting comprehensive sets of molecular concepts (i.e. gene ontologies, pathways, interactions, and domains, etc.) to compute the concept signatures shared by the highly weighted genes, then testing the pathway enrichments over all genes sorted by their functional similarity to the highly weighted genes based on their associations with these concept signatures. WCSEA takes a broader approach for assessing gene set relations and leverages universal concept signature to tolerate the stochasticity of the data. The utilization of the universal concept signature for PE analysis will be particularly useful for scRNA-seq data as technical noise and dropouts in gene expressions are less likely to affect the universal concept signature. This creates a “buffer zone” to tolerate the technical noise of SCS data and improves reproducibility through leveraging the stable concept signatures of the highly weighted genes for PE analysis.

Through analysis of three single cell transcriptomics datasets, we demonstrate the outstanding performance and reproducibility of CSEA and WCSEA on detecting complex pathway alterations under high noise and low coverage of scRNAseq data. Under simulated technical variability of scRNA-seq, CSEA and WCSEA yielded lowest deviations in PE results compared to other commonly used PE methods. More importantly, we demonstrate the outstanding stability of IndepthPathway results in a real dataset of matched scRNAseq data and bulk RNAseq data for the same comparing cell line models. Whereas existing tools such as GSEA, NEA, ORA, iDEA, and UniPath, showed much larger deviations. This is consistent with the report by Holland, C.H. et al (Holland, et al., 2020), that the performance of all PE tools they tested dropped significantly with decreasing gene coverage. They suggest that applying PE tools on perturbation-based pathway gene sets which they termed as footprint gene sets, are more effective. Such approach, however, will leave away most function-based pathway gene sets and limit the analysis to perturbation-based gene sets. WCSEA can endure the stochasticity of single cell sequencing data and detect the enrichment of functional-based pathways with high reproducibility, thus substantially improving the scientific rigor of the pathway analysis for single cell groups. Furthermore, we provided five modules for visualization of the pathway enrichment results. In particular, the pathway association heatmap and network allows for visualizing the functional associations of top enriched pathways to reveal the pathway functional clusters (**Figure 2A**). The visualization of differential gene expression in the context of interactome network allow for exploration of functional links between the gene of interest and the selected top pathways. This function is particularly helpful for genetic perturbation studies to examine the interactions between the perturbated gene of interest and the up or downregulated pathways (**Supplementary Fig. 5**). The development of the IndepthPathway tool will promote the application of single cell sequencing technology to explore the cellular pathway mechanisms at more precise resolution.

Merging a wide array of concept datasets carries inherent concept redundancy, as different gene set sources follow different nomenclatures, and different categories of concepts possess different levels of inherent redundancy. Our uniConSig algorithm possesses a unique ability to compute the redundancy within molecular concept data and sieve only the unique parts of signature concepts for further calculation via an innovative mathematical scheme to penalize the partially overlapping concepts (**Fig. 1B**). This allows for comprehensively assessing the function relations through merging a wide array of molecular concept datasets. The new design of Weighted CSEA (WCSEA) algorithm will empower PE analysis based on a weighted gene list to provide more power to detect different magnitudes of pathway alterations reflected by the levels of differential expressions and better tolerance of the technical noise of scRNA-seq. WCSEA and GSEA has similar functionality by intaking a weighted gene list and producing pathway enrichment results, under different designs. Based on the strength of WCSEA in inferring the functional enrichments of gene sets, and the strength of GSEA in direct measuring the enrichments of gene sets, we recommend the use of WCSEA for interpretation of functional-based pathways and the use GSEA for exploring the enrichment of perturbation-based gene expression signature.

Purpose-designed PE methods such as IndepthPathway that can better address the characteristics of scRNA-seq data will be of utmost importance to ensure the accuracy and reproducibility of the PE results. The integration of the up and downstream analysis modules such as differential gene expression, pathway disambiguation, functional associations of pathways, and heatmap and network visualization of the top enriched pathways will greatly enhance the functionalities of IndepthPathway.

## Author contributions

S. L. performed the analyses for the three scRNAseq datasets, carried out the simulation studies, and cowrote the manuscript. L.D. prepared the first Github submission and contributed to parts of Figure 1–2. Y. W. performed network analysis. M.A.S. and K.W. advised on the biostatistics and the R package for public release. X-S. W. designed the study, developed the IndepthPathway package, performed PE analysis on the HSC dataset, and wrote the manuscript.

## Code availability

The R modules for IndepthPathway are available through: https://github.com/wangxlab/IndepthPathway

## Acknowledgements

This study was supported by P30 CA047904 CCSG Dev Funds (X-S. W.), NIH grant 1R01CA181368 (X-S.W.), and 1R01CA183976 (X-S.W.). This study is also in part supported by 1R21CA237964 (X-S.W.), P50 CA097190 Head and Neck SPORE DRP (X-S. W.), PA breast cancer coalition (X-S.W.), Commonwealth of PA Tobacco Phase 15 Formula Fund (X-S. W.), the Shear Family Foundation, and the Hillman Foundation. We thank Yuehua Zhu and Megan E. Yates for testing the IndepthPathway package. This research was supported in part by the University of Pittsburgh Center for Research Computing through the resources provided. We especially thank Fangping Mu for his kind assistance with the research computing. This work also used the Extreme Science and Engineering Discovery Environment (XSEDE) supported by National Science Foundation grant number OCI-1053575. Specifically, it used the Bridges system, which is supported by NSF award number ACI-1445606, at the Pittsburgh Supercomputing Center (PSC).

