## Supplementary Materials for "IndepthPathway: an integrated tool for in-depth pathway enrichment analysis based on bulk and single cell sequencing data"

**SUPPLEMENTARY FIGURES**

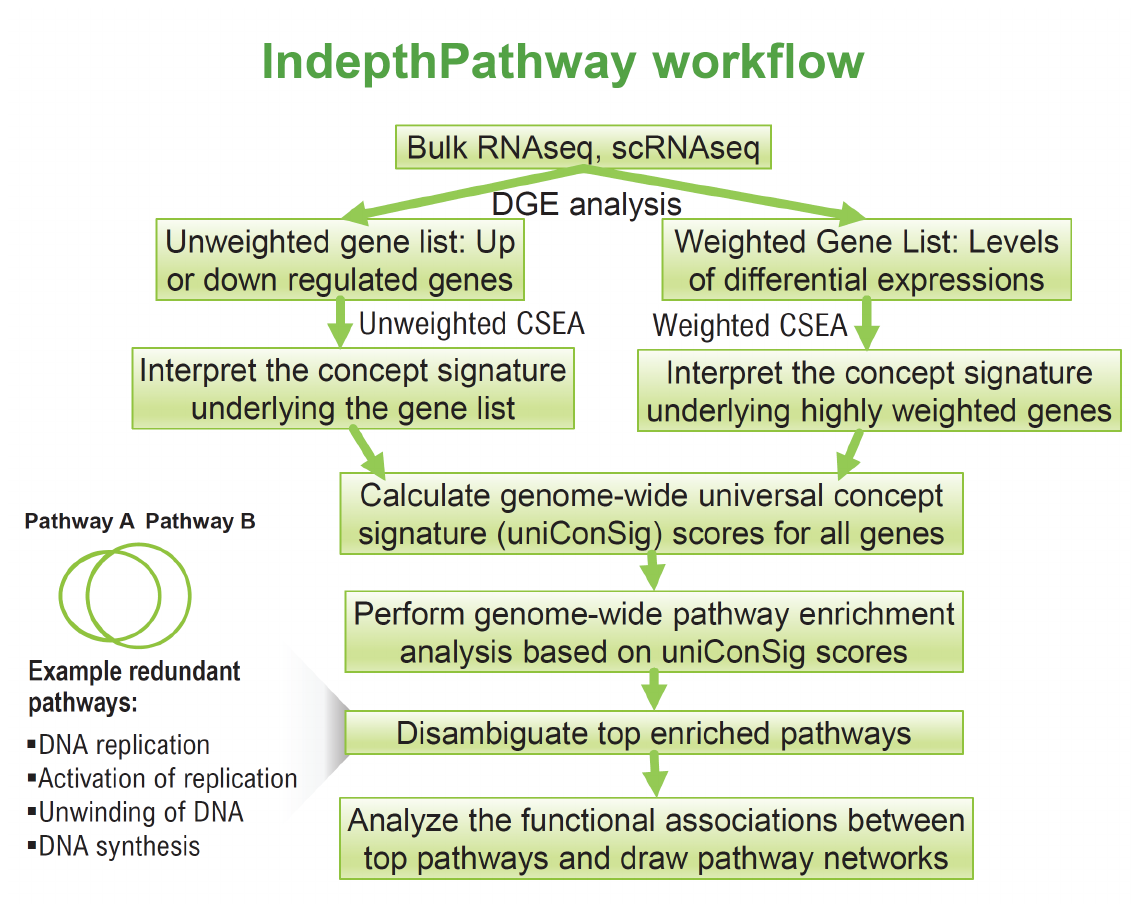

**Supplementary Figure 1.** The workflow of the IndepthPathway package. The diagram depicts the difference between CSEA and WCSEA. CSEA and WCSEA have distinct functionality and accept different types of inputs for the PE analysis. CSEA accepts a list of target genes (i.e., up, or downregulated genes), whereas WCSEA accepts a weighted target gene list (i.e., levels of differential expressions) as input data. CSEA interprets pathways in a target gene set, whereas WCSEA interprets pathways in a weighted gene list.

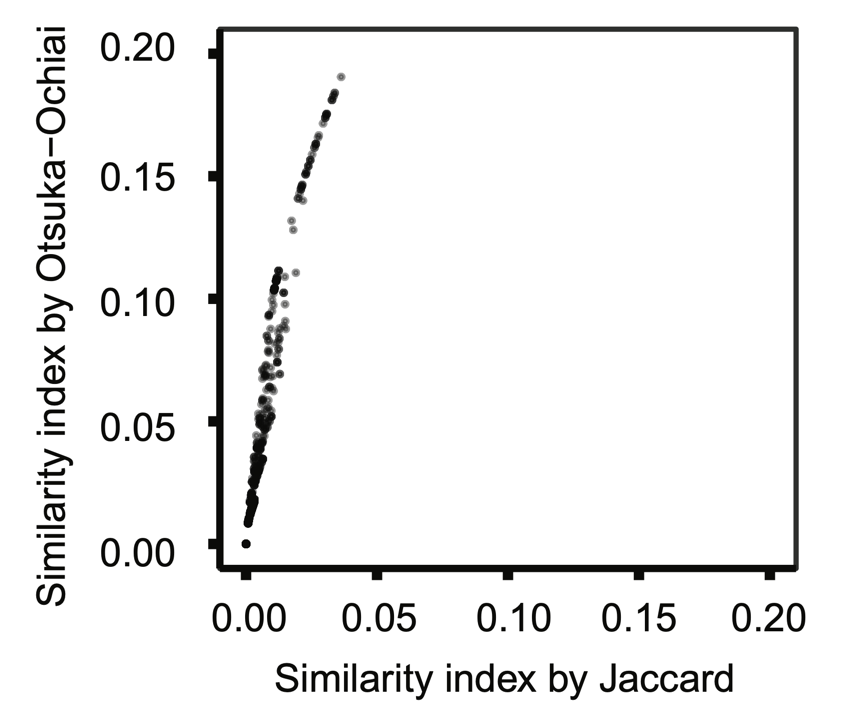

**Supplementary Figure 2.** Comparing the performance between Otsuka-Ochiai coefficient and Jaccard Index for estimating the similarities between overlapping molecular concepts of large and small sizes. We identified 300 large and small concept pairs in which each large concept contains all genes in the paired small concepts, and calculated similarity indexes by both methods. X-axis is the similarity index by Jaccard method and y-axis is the similarity index by Otsuka-Ochiai method. The similarity indices by Otsuka-Ochiai method are larger than the ones by Jaccard method thus better reflect the similarities between these overlapping concepts.

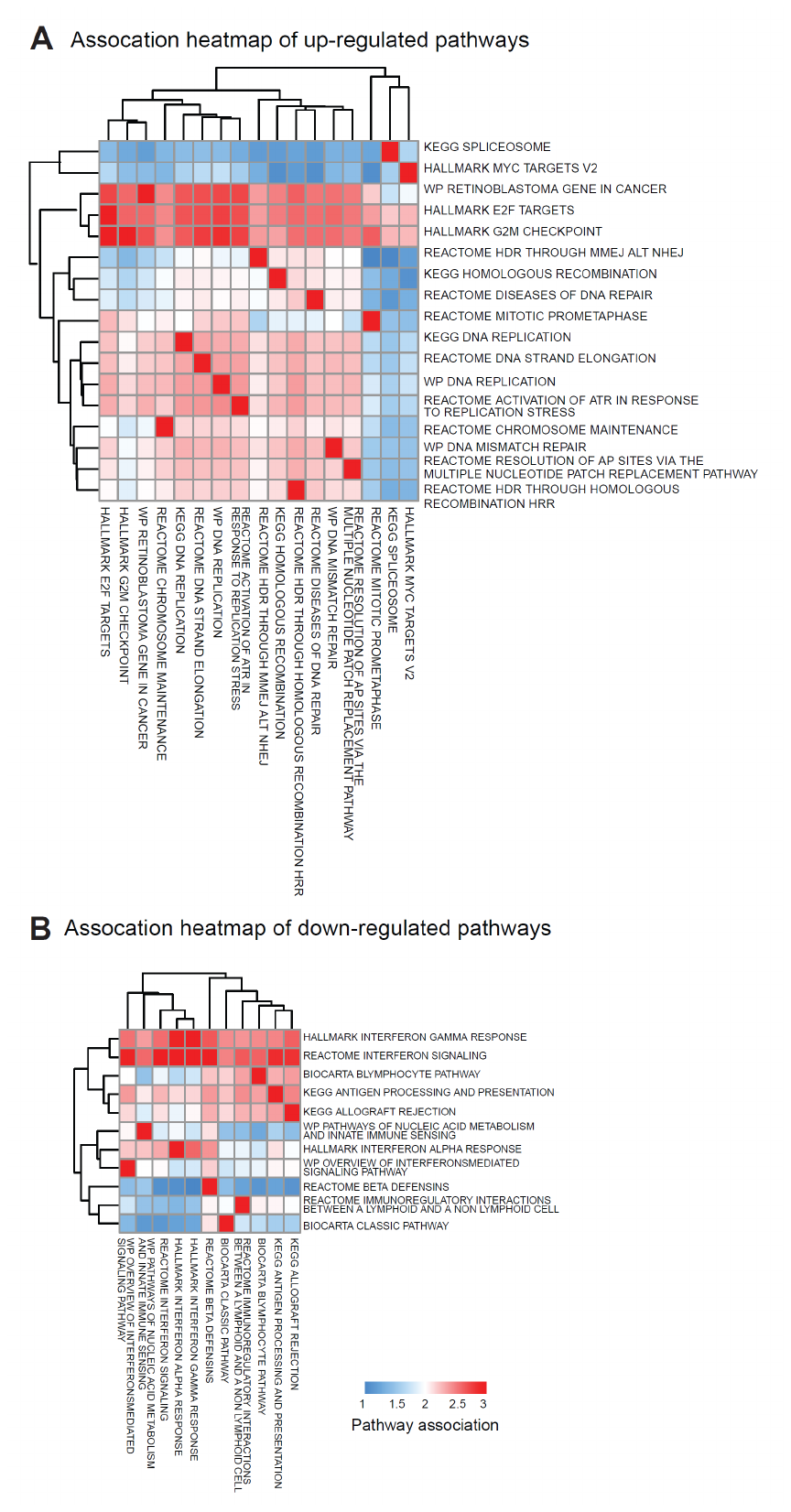

**Supplementary Figure 3.** Heatmap visualizing the functional similarity between top enriched pathways from WCSEA analysis. The PE analysis was performed to compare active and quiescent HSC cells based on the HSC scRNA-seq dataset (GSE69981). The heatmaps show the functional associations between top upregulated pathways (A) or downregulated pathways (B).

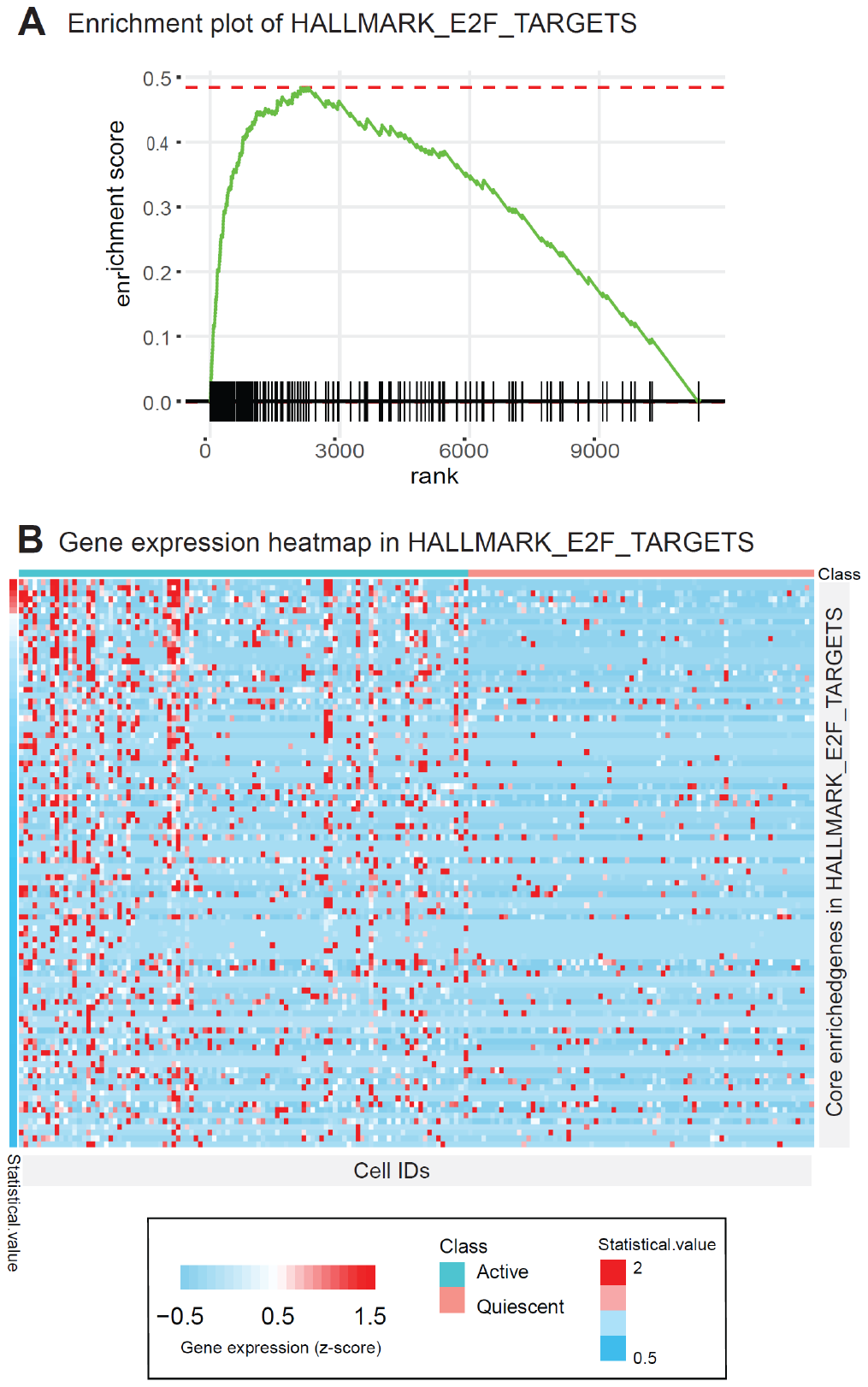

**Supplementary Figure 4.** Visualizing the enrichment of a selected pathway in the weighted gene list reflecting the differential expression levels between active and quiescent HSC cells. (A) The enrichment plot of “HALLMARK_E2F_TARGETS” pathway. (B) Heatmap showing core enriched genes of the HALLMARK_E2F_TARGETS pathway.

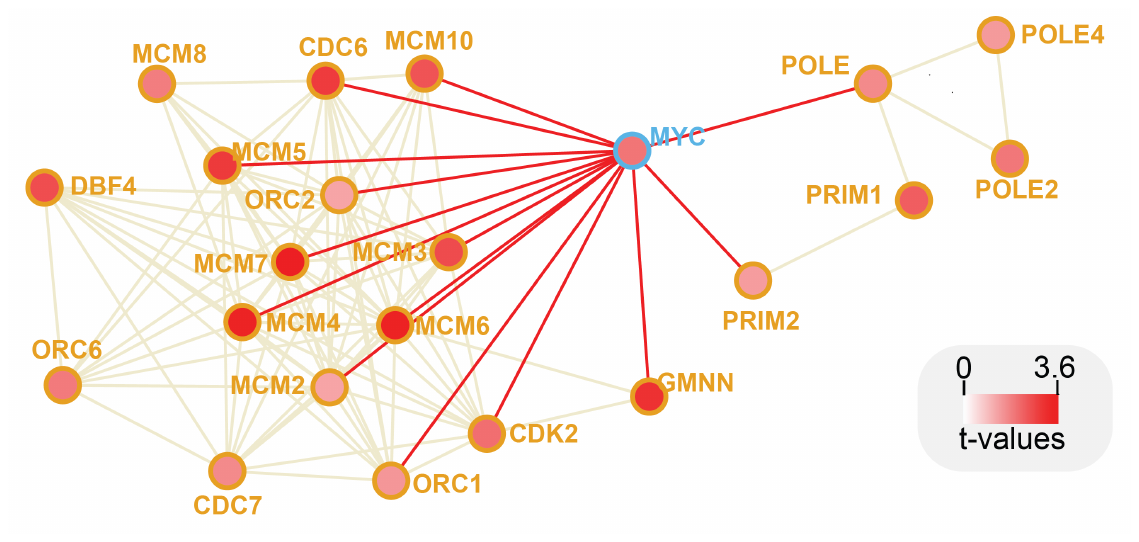

**Supplementary Figure 5**. Interactome network of MYC and the selected pathway “activation of the replicative complex" (REACTOME) in the context of differential expression during quiescent to active cell state transition.

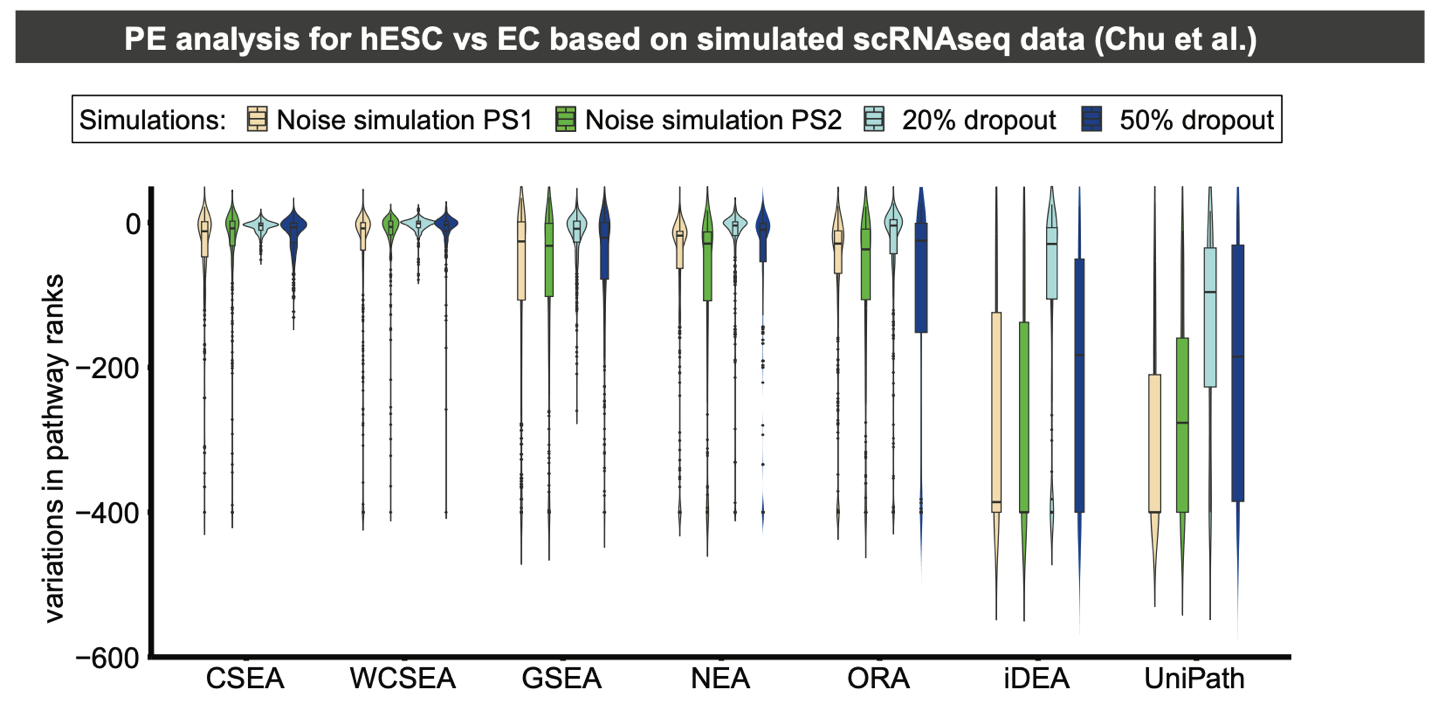

**Supplementary Figure 6.** The reproducibility of pathway enrichment methods under simulations of scRNA-seq technical variability and dropouts in gene expression in the hESC dataset. Noise simulations are performed using SymSim with two sets of parameters representing different noise levels based on the bulk RNAseq data for EC and hESC. The dropouts are simulated via randomly deleting 20% or 50% expressed genes. Read counts from the original bulk RNAseq are used as true counts before simulations. SCDE is used for DE analysis, and the results are used for PE analysis. Variations in pathway ranks for top 30 up- or down-regulated pathways are used for benchmarking with negative variations indicating reduced significance of the pathways.

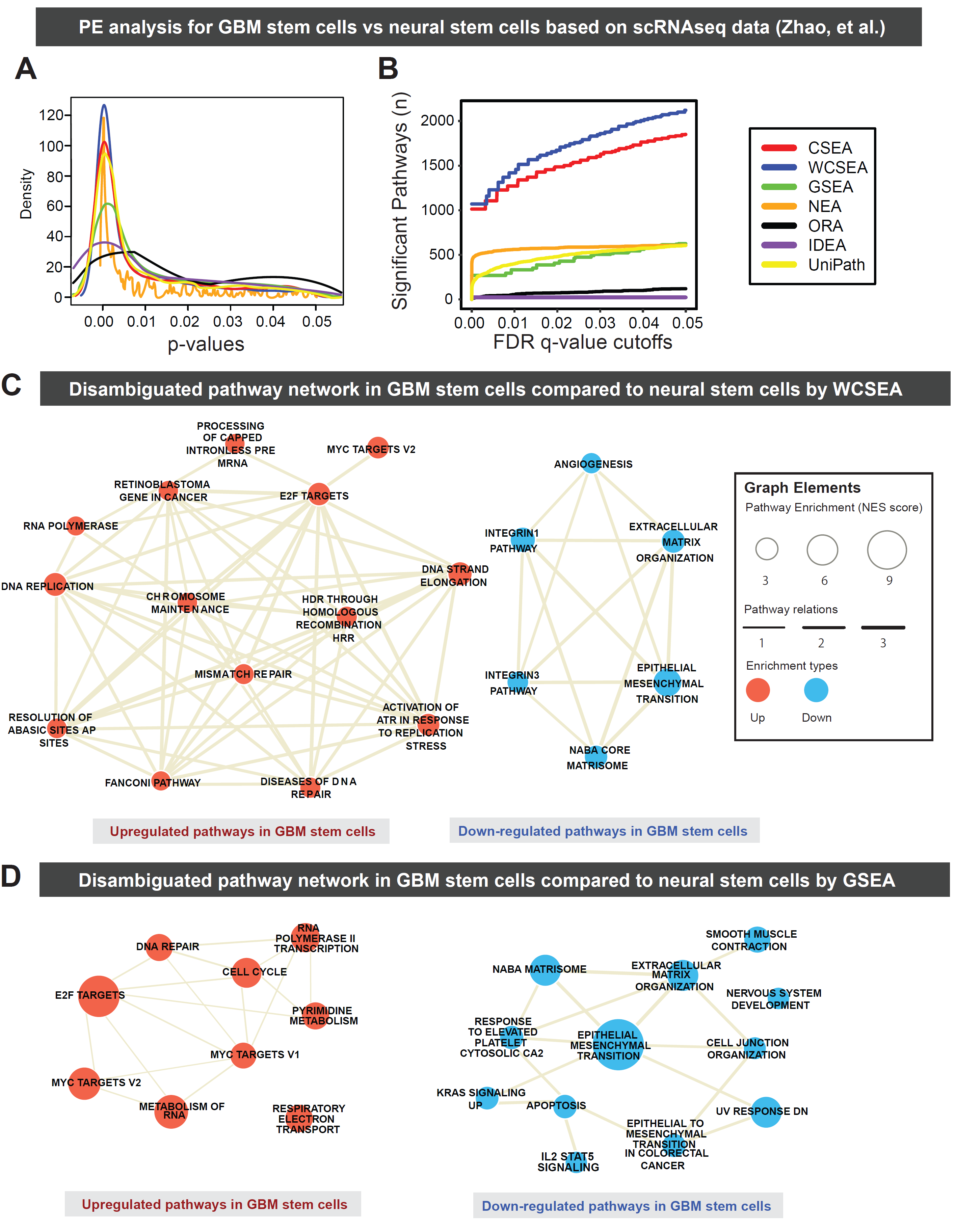

**Supplementary Figure 7.** Pathway enrichment results from different PE tools comparing neural stem cells with GBM stem cells. (**A**) Kernal density for pathway p-values from CSEA, WCSEA, and other PE tools. (**B**) Number of significantly enriched pathways detected by PE tools based on the increasing FDR q-value cutoffs from 1×10^-05^ up to 0.05. (**C-D**) Pathway association networks showing the top up-regulated and down-regulated pathways in GBM stem cells compared to neural stem cells detected by WCSEA (**C**) or GSEA (**D**).

**Pseudo-code of WCSEA**

\#### ======== Step1. Call necessary databases ======== ####

\#### ======== #### ======== #### ======== #### ======== ###

\# Call the modules that are necessary to run uniConSig, WCSEA, and make heatmap, network, and enrichment plots.

> CALL indepthPathway modules

\# Molecular concepts were compiled from Molecular Signatures Database (MSigDB) Hallmark, C2CP pathways (v.7.0),

\# NCBI EntrezGene interactome and conserved domain database

> CALL precompiled molecular concept database

\# This preCal matrix will be used to penalize the redundancy between molecular signatures associated with each gene.

\# This will reduce the inflation effect on the uniConSig scores.

> CALL preCal matrix

\# The Hallmark and C2CP pathway are provided for WCSEA pathway enrichment analysis. Users can use C2 Reactome,

\# C5 Gene Ontology Biolgical Pathway, or any other pathway collections

> CALL MSigDB Hallmark and C2CP pathway data

\#### ======== Step2. Read gene expression data and run SCDE or Limma ======== ####

\#### ======== #### ======== #### ======== #### ======== ### ======== ######

\# A user should make gct and cls files in advance. The class has two phenotype elements such as quiscent vs active,

\# or tumor vs normal

> INPUT Gene expression gct and class cls files

\# SCDE or Limma will identify Up- or down-regulated DEGs which will be used as target genes.

> RUN SCDE or Limma to identify differentially expressed genes.

\# The 'signed q-values' are calculated for the target genes. The values become gene weights

> CALCULATE signed q-value for the target genes

\#### ========	Step3. weighted K-S test and uniConSig	======== ########

\#### ======== #### ======== #### ======== #### ======== ### ======== #####

\# Weighted K-S test will find enriched molecular concepts. Up-regulated DEGs will have positive weights and donw-regulated

\# DEGs will have negative weights.

> RUN weighted K-S test for the genes against molecular concept signatures

\# This will calculate up.uniConSig and down.uniConSig scores of genes. uniConSig scores will represent gene functions

\# underlying biological or pathological processes

> CALCULATE uniConSig scores for wide quantification of human gene functions

\#### ======== Step4. WCSEA for pathway enrichment	======== ##########

\#### ======== #### ======== #### ======== #### ======== ### ======== ###

\# This will identify up-regulated pathways in a certain class

> INPUT up.uniConSig scores of genes to WCSEA

> RUN WCSEA to perfrom deep functional assessment for pathway enrichment

\# This will identify down-regulated pathways in a certain class

> INPUT down.uniConSig scores of genes to WCSEA

> RUN WCSEA to perfrom deep functional assessment for pathway depletion

\#### ======== Step5. Deambiguation to correct crosstalk effect ======== ##########

\#### ======= #### ======= #### ======= #### ======= ### ======= ##### ====== ##

\# Users can select top 30 up-regulated pathways

> SELECT Top 30 up-regulated pathways

> Up.DEAMBIGUATE the WCSEA results to identify independent pathways in up.wCSEA.result

\# Users can select top 30 down-regulated pathways

> SELECT Top 30 down-regulated pathways

> Down.DEAMBIGUATE the WCSEA results to identify independent pathways in down.wCSEA.result

\#### ======== Step6. Visualization for the WCSEA results	======== ##########

############# ========== ############# ========== ############# =======

\# Users can make 5 different visualization to represent WCSEA results

\# VISUALIZE_Option1 is to make association heatmap to represent the functional association between selected pathways

> SELECT Top 30 up-regulated pathways from up.deambiguated pathways

> CALCULATE and DRAW association heatmap for the selected pathways

> SELECT Top 30 down-regulated pathways from down.deambiguated pathways

> CALCULATE and DRAW association heatmap for the selected pathways

\# VISUALIZE_Option2 is to make network to represent the functional association between selected pathways

> SELECT Top 30 up-regulated pathways

> SELECT Top 30 down-regulated pathways

> MERGE the up- and down-regulated pathways

> DRAW network for the merged pathways

\# VISUALIZE_Option3 is to make enrichment plot for a selected pathway.

> SELECT a pathway and DRAW enrichment plot.

\# VISUALIZE_Option4 is to make heatmap of gene expression data for a selected pathway

> SELECT a pathway

> INPUT Gene expression gct and class cls files

> SUBSET the gene expression data by the genes that belong to a selected pathway

> DRAW heatmap of gene expression subset data

\# VISUALIZE_Option5 is to make interactome network for a target gene of user’s interest and the selected pathways.

> INPUT interactome data

> GET the target gene list of user interest

> GET weight for the target gene from SCDE or Limma result

> DRAW interactome network for the target genes
